# Generating an organ-deficient animal model using a multi-targeted CRISPR-Cas9 system

**DOI:** 10.1101/2023.07.31.551401

**Authors:** Jonathan Jun-Yong Lim, Shunsuke Yuri, Taro Kawai, Ayako Isotani

## Abstract

Gene-knockout animal models with organ-deficient phenotypes used for blastocyst complementation are generally not viable. Animals need to be maintained as heterozygous mutants, and homozygous mutant embryos yield only one-fourth of all embryos. In this study, we generated organ-deficient embryos using the CRISPR-Cas9-sgRNA^ms^ system that induces cell death with a single-guide RNA (sgRNA^ms^) targeting multiple sites in the genome. The Cas9-sgRNA^ms^ system interrupted cell proliferation and induced cell ablation *in vitro*. The mouse model had Cas9 driven by the *Foxn1* promoter with a ubiquitous expression cassette of sgRNA^ms^ at the *Rosa26* locus (*Foxn1^Cas^*^9^; *Rosa26_ms*). It showed an athymic phenotype similar to that of nude mice but was not hairless. Eventually, a rat cell-derived thymus in an interspecific chimera was generated by blastocyst complementation of *Foxn1^Cas^*^9^; *Rosa26_ms* mouse embryos with rat embryonic stem cells. Theoretically, half of the total embryos have the Cas9-sgRNA^ms^ system because Rosa26_ms could be maintained as homozygous.

## Introduction

With increase in the demand for organ transplants in recent decades, organ shortage due to insufficient number of organ donors has become a worldwide problem. This deficit may be compensated by growing transplantable human organs in laboratory animals using the blastocyst complementation method. Injecting pluripotent stem cells (PSCs) into organ-deficient embryos can regenerate fully developed organs^1–5^. Organ-deficient embryos are hence necessary for blastocyst complementation.

Current methods for generating organ-deficient embryos include knockout (KO) of organ-specific genes^1–8^ and conditional cell ablation with diphtheria toxin A (DTA) using the Cre/LoxP system^9–12^. However, most organ-deficient animal models have a lethal phenotype. Therefore, animals need to be maintained as heterozygous mutants or in two or more lines, and organ-deficient embryos yield only a quarter or less of the total number of embryos in most crossing combinations. Thus, using gene-KO and DTA-based cell ablation models for blastocyst complementation is labor-efficient and costly. Most genes that control organ and tissue development are also involved in other processes. The injection of PSCs rescued KO cells in some gene-KO models, resulting in organs containing a mixture of host-and PSC-derived tissues.

Clustered regularly interspaced palindromic repeats (CRISPR)-Cas9 technology has revolutionized gene editing, whereby DNA double-strand breaks (DSB) can be induced at any desired site in the genome, corresponding to customized single-guide RNA (sgRNAs)^13^. CRISPR-Cas9 is typically used to induce DNA DSBs at single sites for gene editing. Nevertheless, it is hypothesized that programmed cell death can be triggered in cells by inducing multiple DNA DSBs. This technology can kill cancer cells by inducing multiple DNA DSBs in their genome^14, 15^ or by targeting cancer-specific fusion oncogenes^16^ without affecting healthy cells. However, this method has not been experimented to induce cell ablation in organ-specific cells to produce organ-deficient animal models. Hence, we aimed to investigate whether organ-deficient animal models could be produced using multiple-target CRISPR-Cas9 (Figure 1A).

**Figure 1.**
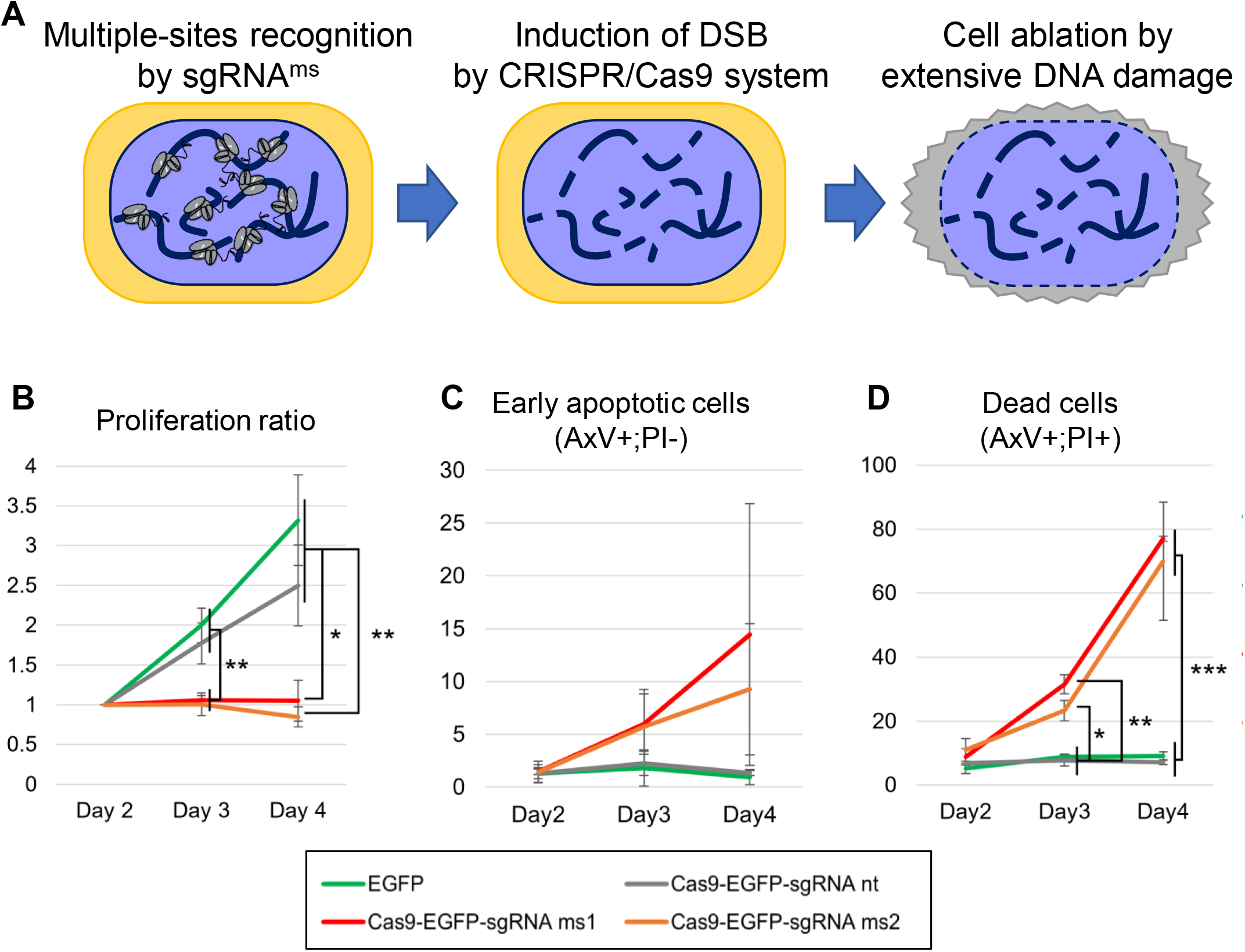
Cell ablation potency of the Cas9-sgRNA^ms^ system in HEK293T. (A) The strategy of the Cas9-sgRNA^ms^ system is to induce cell ablation through multiple-site DNA double-strand breaks (DSBs). (B) Cell proliferation after transiently expressing Cas9-sgRNA^ms^ in HEK293T cells. The value of the Y-axis indicates the relative number of GFP-positive cells. (C) Relative levels of early apoptotic cells after transient expression of Cas9-sgRNA^ms^ in HEK293T cells. The value of the Y-axis indicates the relative cell population of AxV+;PI-;GFP+ cell (%) / GFP+ cell (%). (D) Relative levels of dead cells after transient expression of Cas9-sgRNA^ms^ in HEK293T cells. The value of the Y-axis indicates the relative cell population of AxV+;PI+;GFP+ cell (%) / GFP+ cell (%). Cas9-sgRNA^ms^ transfected cells were monitored using EGFP signals. Cas9-sgRNA^nt^ has an empty target recognition sequence in the sgRNA and was used as a control in (B)–(D). One-way ANOVA, Tukey honestly significant difference test was used in Figures 1B to 1D. *p < 0.05; **p < 0.01; ***p < 0.001.

The thymus is the primary organ of the adaptive immune system and is responsible for the education, maturation, and differentiation of immune cells. These immune cells recognize self-and non-self-antigens and are responsible for solid graft rejection in organ transplants^17^. Thymus organogenesis and thymic epithelial cell differentiation are controlled by a single master regulator *Foxn1*^18^ that serves as an ideal candidate for this proof-of-concept study. The athymic, T cell-deficient, viable *Foxn1^-/-^* mouse model^19–21^ can generate a rat PSC-derived thymus via blastocyst complementation^2^. Therefore, we attempted to generate a thymus-deficient mouse model by introducing multiple DSB into Foxn1-expressing cells in this study.

## Results

### The Cas9-sgRNA^ms^ system affected cell proliferation and induced apoptosis in HEK293T cells

Two multiple-site-targeting sgRNAs^ms^ were designed consisting of repeating nucleotide sequences to recognize the Cas9 genome (Table 1). HEK293T cells were co-transfected with the Cas9-sgRNA^ms^-expressing plasmid and the corresponding target DNA sequence of sgRNA^ms^ in the pCAG-EGxxFP plasmid to validate the cleavage activity and target specificity of the Cas9-sgRNAs^ms^ system. EGFP expression indicated the successful cleavage of the target DNA sequence^22^. An EGFP signal was observed in most cells transfected with Cas9-sgRNA^ms1^ or Cas9-sgRNA^ms2^ and pCAG-EGxxFP with the target DNA sequence (Figure S1). Minimal EGFP signals were detected when sgRNA^ms1^ was used to cleave the ms2 target sequence in pCAG-EGxxFP and vice versa. This further indicated that the targeting of the two sgRNA^ms^ was unique.

**Table 1.**
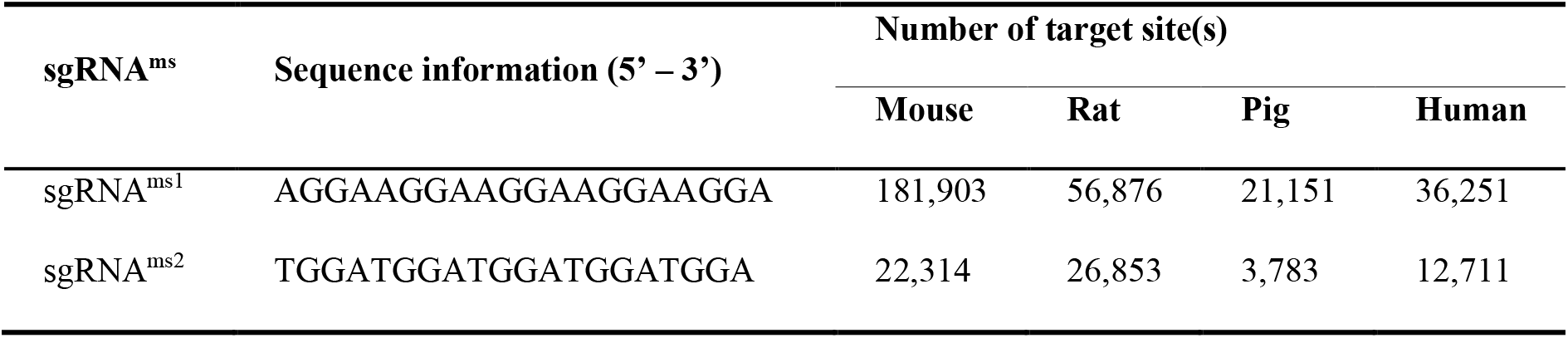
Sequence of sgRNA^ms^ and number of target sites.

Furthermore, Cas9-T2A-EGFP was transiently expressed with or without sgRNA^ms^ in HEK293T cells. There was a lag in cell growth in the presence of Cas9-sgRNA^ms^ compared to that in the non-targeting (nt) sgRNA control (Figure 1B). The cells were stained with annexin V (AxV) and propidium iodide (PI) and analyzed using flow cytometry to further understand whether the cells were undergoing apoptosis and cell death. Early apoptotic cells were defined as AxV^+^ and PI^-^ cells, whereas dead cells were defined as AxV^+^ and PI^+^ cells. Gated EGFP^+^ cells (Cas9-expressed cells) transfected with sgRNA^ms1^ or sgRNA^ms2^ showed a significant increase in dead cell population between days 2 and 4 post-transfection (Figures 1D). Early apoptotic cell population tended to increase, but no significant differences were observed (Figures 1C).

### The Cas9-sgRNA^ms^ system induced multiple DNA double-strand breaks in HEK293T cells

Doxycycline-inducible Cas9-T2A-EGFP in HEK293T cell lines that constitutively express sgRNA^ms^ were established to understand the downstream effects of the Cas9-sgRNA^ms^ system and minimize the variation in transfection efficiency between cells (Figure S2). Two cell lines for each sgRNA^ms^ were used in all subsequent experiments. We investigated the DNA DSB, apoptosis, and cell proliferation after 72 h of doxycycline-induced Cas9 expression. Histone H2AX phosphorylated at serine 139 (γH2AX) accumulates at DNA damage sites and serves as a molecular marker of DNA DSBs^23^. We stained doxycycline-induced cells for γH2AX to detect DNA DSBs as our Cas9-sgRNA^ms^ system was designed to cleave DNA at multiple sites. Cells that expressed sgRNA^ms^ showed a significant increase in γH2AX signal intensity compared to cells expressing non-targeting (nt) sgRNA as a control (Figures 2A and B). Furthermore, we observed a significant increase in the percentage of early apoptotic cells (AxV^+^;PI^-^) and dead cells (AxV^+^;PI^+^) compared to that in the control (Figures 2C), albeit at low levels. This suggests that the Cas9-sgRNA^ms^ system altered cell growth through DNA DSB but did not induce apoptosis within this time frame.

**Figure 2.**
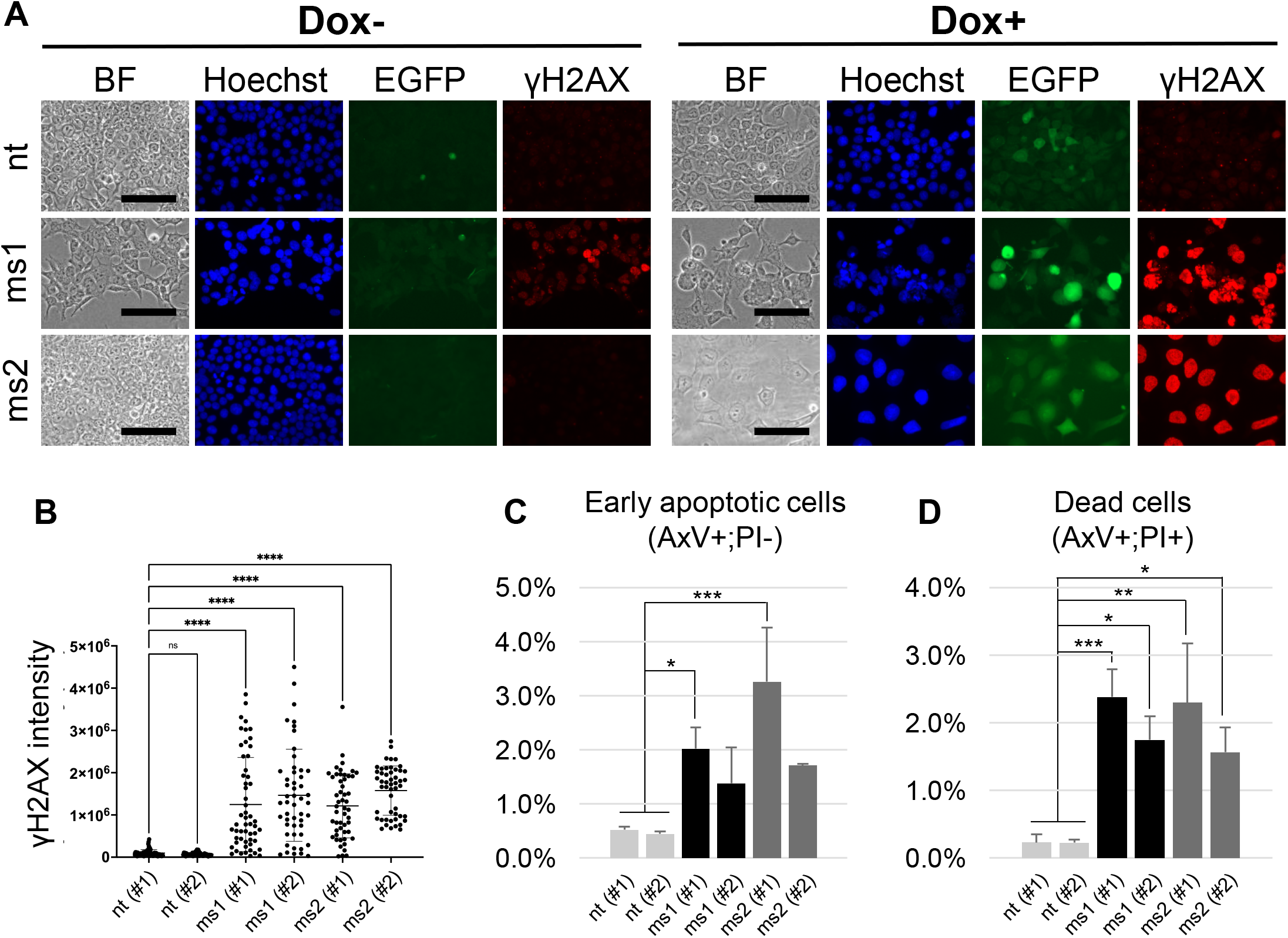
Induction of DNA DSB in HEK293T by the Cas9-sgRNA^ms^ system. (A) DNA DSB assays detected γH2AX accumulation. sgRNA^ms^ were driven by the ubiquitously expressing U6-promoter, and Cas9 expression was induced by Doxycycline; the Cas9-sgRNA^ms^ system functioned after adding doxycycline to the culture medium. BF means bright field. Blue: nuclei were stained by Hoechst33342, green: Cas9 expression was monitored by EGFP signal, and red: DNA DSBs were indicated by γH2AX accumulation. Images show data from the nt (#2), ms1 (#2), and ms2 (#2) cell lines. The other cell lines showed the same results. The data were quantitatively analyzed as shown in Figure 2B. (B) Quantitative analysis of γH2AX intensity. (C) The rate of early apoptotic and (D) dead cells. One-way ANOVA, Tukey honestly significant difference test was used in Figures 2B, 2C and 2D. *p < 0.05; **p < 0.01; ***p < 0.001; ****p < 0.0001. “ns” in Figure 2B means not significant differences.

The cell proliferation rate of cells significantly decreased with the expression of sgRNA^ms1^ and sgRNA^ms2^ compared to the controls using an MTT [3-(4,5-dimethylthiazol-2-yl)-2,5-diphenyltetrazolium bromide] assay to compare cell proliferation in the presence and absence of doxycycline (Figure S3). The decrease in cell proliferation was negatively affected. Therefore, we hypothesized that DNA synthesis might also be affected by the Cas9-sgRNA^ms^ system. BrdU is incorporated into DNA during DNA synthesis in cells, we performed BrdU staining and showed that there was a significant increase in the proportion of BrdU-negative cells among cells expressing sgRNA from a minimum of 100 cells counted for each cell line (Figure 3). The nuclei in Cas9-sgRNA^ms^ modified cells were larger than those in the control. Taken together, these results suggest that the designed Cas9-sgRNA^ms^ system induced multiple DNA DSBs throughout the genome and negatively affected HEK293T cell proliferation.

**Figure 3.**
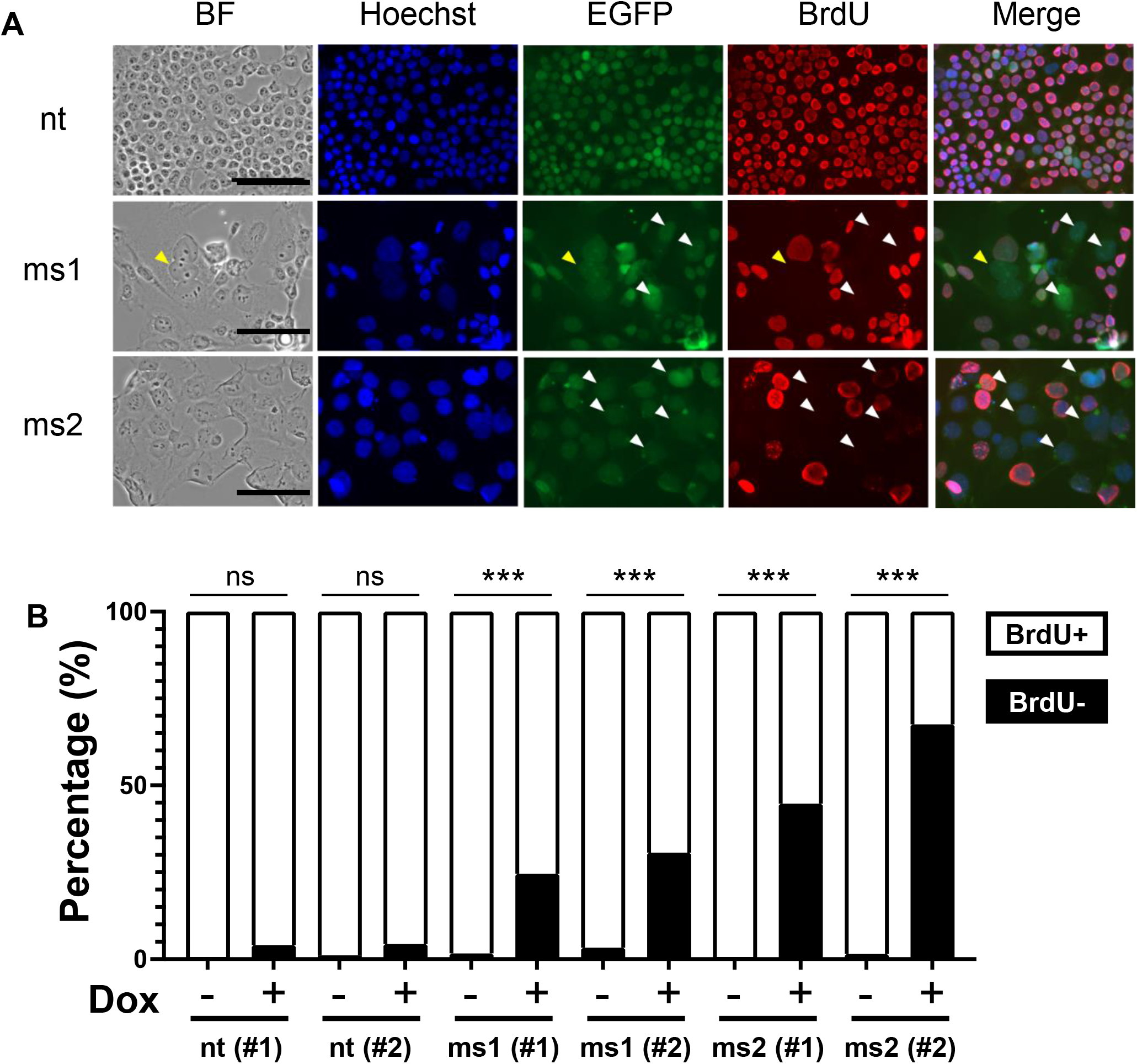
Obstruction of HEK293T cell proliferation by the Cas9-sgRNA^ms^ system. (A) Cell proliferation assay with BrdU staining. BF indicates bright field. Blue: nuclei were stained by Hoechst33342, green: Cas9 expression was monitored by EGFP signal, and red: DNA synthesis was indicated by BrdU incorporation. Pictures show data from the nt (#2), ms1 (#2), and ms2 (#2) cell lines treated with Dox. The other cell lines showed the same results. The data were quantitatively analyzed as shown in Figure 3B. (B) Quantitative analysis of DNA synthesis via BrdU incorporation. Fisher’s exact probability test was used in (B). *** P < 0.001. “ns” in Figure 3B means no significant differences.

### The Cas9-sgRNA^ms^ system induced cell ablation in mESCs

The aim of this study was to ablate organ-specific cells in mouse models. Therefore, it was necessary to further test whether the Cas9-sgRNA^ms^ system was feasible in mouse cells. The function of mouse ESC lines with sgRNA^ms^ knocked-in the *Rosa26* locus (*Rosa26_ms*) was tested by transfecting Cas9 and a puromycin-resistant gene-expressing plasmid that has no target sequence of sgRNA in PX459. Fewer ALP^+^ cells were detected puromycin selection compared with wild-type mESCs (Figure S4). This indicated that the simultaneous expression of Cas9 and sgRNA^ms^ induced cell death in most mESCs. The *p53* tumor suppressor gene is known to play a pivotal role in controlling cell cycle arrest and cell death. Therefore, we further investigated whether cell ablation triggered by Cas9-sgRNA^ms^ was related to the *p53* pathway. There was a significant decrease in the relative number of early apoptotic cells (AxV^+^;PI^-^) and dead cells (AxV^+^;PI^+^) when we transfected Cas9-sgRNA^ms^ into two individual *p53*-KO mESC lines (Figure S5). These data suggest that Cas9-sgRNA^ms^-induced cell ablation was related to the *p53* pathway.

### *Foxn1^Cas^*^9^; *Rosa26_ms* mice were athymic and lack peripheral T cells

The thymus was selected as the target organ to generate an organ-deficient mouse model using the Cas9-sgRNA^ms^ system. Thymus-deficient mice were reported to be viable until the adult stage, have immunodeficiency owing to a lack of thymus and peripheral mature T cells, and have disrupted hair growth caused by the *Foxn1* gene^19, 21^. Furthermore, the thymus consists of PSCs formed by blastocyst complementation using *Foxn1^nu/nu^* ^2^. We established founder mouse lines of *Foxn1^Cas^*^9^ and *Rosa26_ms* from mESCs and crossed with wild-type females to obtain F1 progeny respectively. Each F1 mouse was crossed to obtain the next generation. Next generation pups of *Foxn1^Cas^*^9^; *Rosa26_ms* pups were dissected at 9-day postpartum (P9) to examine the thymus phenotype. *Foxn1^Cas^*^9^; *Rosa26_ms* mice were athymic; however, their hair phenotypes were unaffected (Figure 4A).

**Figure 4.**
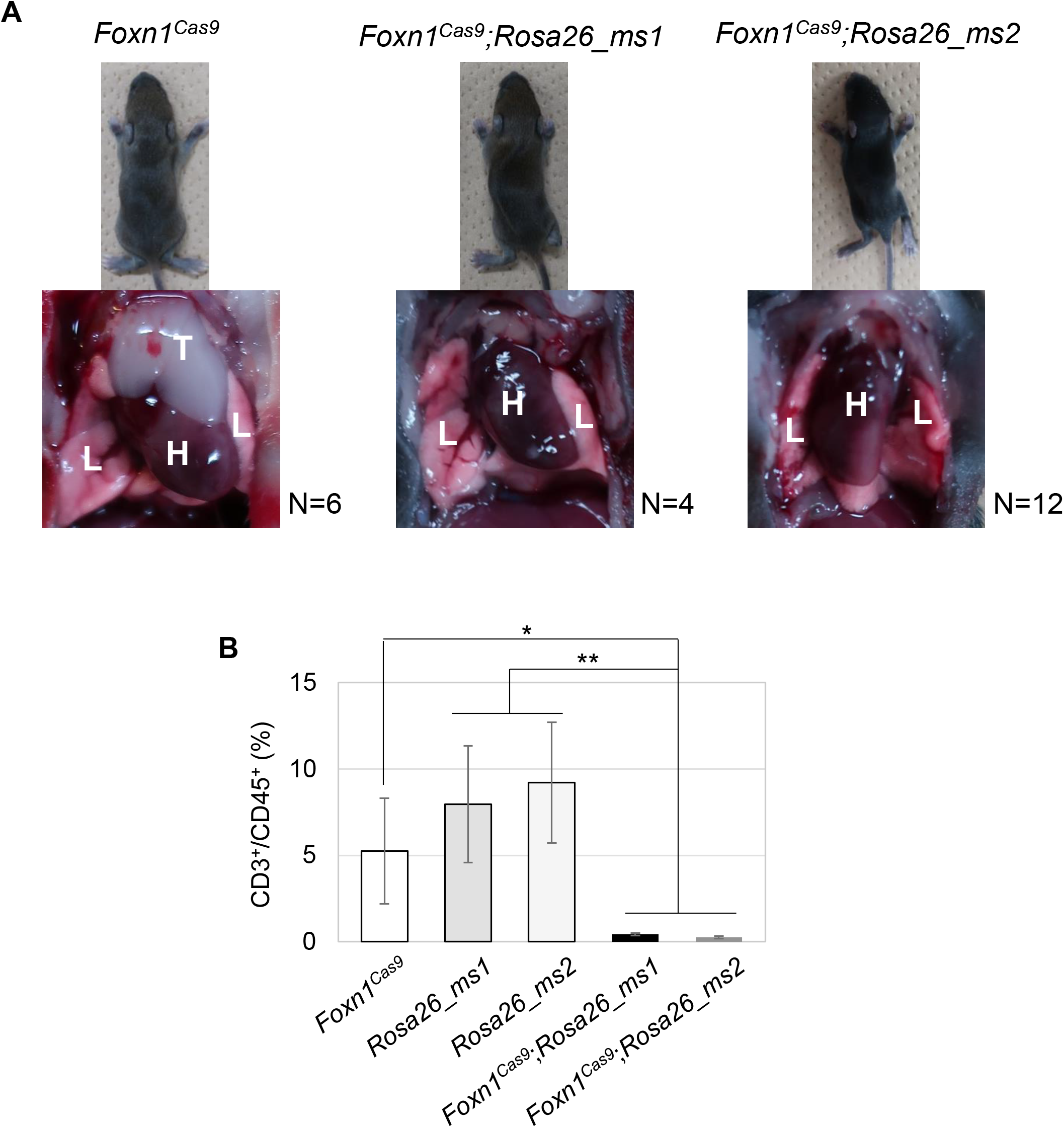
Phenotypes of *Foxn1^Cas^*^9^; *Rosa26_ms* mice. (A) Pictures of nine-day old mice. *Foxn1^Cas9^*; *Rosa26_ms* mice had an athymic phenotype but were unhealthy. *Foxn1^Cas9^*mice were used as normal phenotype controls. Yellow arrows indicate multiple nuclei and white arrowheads indicate Cas9-expressed BrdU negative cells. (B) T cell population in splenocytes determined using flow cytometry. One-way ANOVA, Tukey honestly significant difference test was used in Figure 4B. *p < 0.05; **p < 0.01.

*Foxn1^Cas^*^9^; *Rosa26_ms* mice had significantly fewer peripheral CD3+ and CD45+ cells in the spleen compared to control mice lacking the Cas9-sgRNA^ms^ system according to immunostaining and flow cytometry (Figure 4B). The Cas9-sgRNA^ms^ system thus successfully generated a thymus-deficient mouse model.

### Generation of interspecies chimera in the rat thymus using the Cas9-sgRNA^ms^ system

The rat thymus interspecies chimera was generated using athymic mouse models with the Cas9-sgRNA^ms^ system. Rat ESCs (rESCs) tagged with EGFP were injected into mouse blastocysts that were obtained from *Rosa26_ms* females crossed with *Foxn1^Cas9^*; *Rosa26_ms* males, followed by transfer to pseudo pregnant female mice to continue embryo development (Figure 5A). The thymus from the dissected fetuses of *Foxn1^Cas9^*; *Rosa26_ms* mice at E18.5 had high EGFP expression (Figure 5B). However, the morphology of the complemented thymus was slightly different from that of the control thymus, which was used as a host for the non-working Cas9-sgRNA^ms^ system for genotyping. Therefore, we analyzed the thymus sections to understand the contribution of rESC-derived cells. The thymus of the control host contained partially rat-derived thymic epithelial cells. In contrast, the thymic epithelial cells from the *Foxn1^Cas9-^*; *Rosa26_ms* host were mostly derived from rESCs (Figure 5C). Immunostaining of these thymi with cortex (K8)-and medullary (K5)-specific markers showed that rESCs contributed to the cortical and medullary regions in the thymus (Figure 5C). The cortex marker (K8) used in this study could only detect mouse tissue, not rat tissue (Figures 5C and S6). Taken together, these data suggest that *Foxn1^Cas9^*; *Rosa26_ms* mice create an empty thymus niche that can be complemented with injected rESCs.

**Figure 5.**
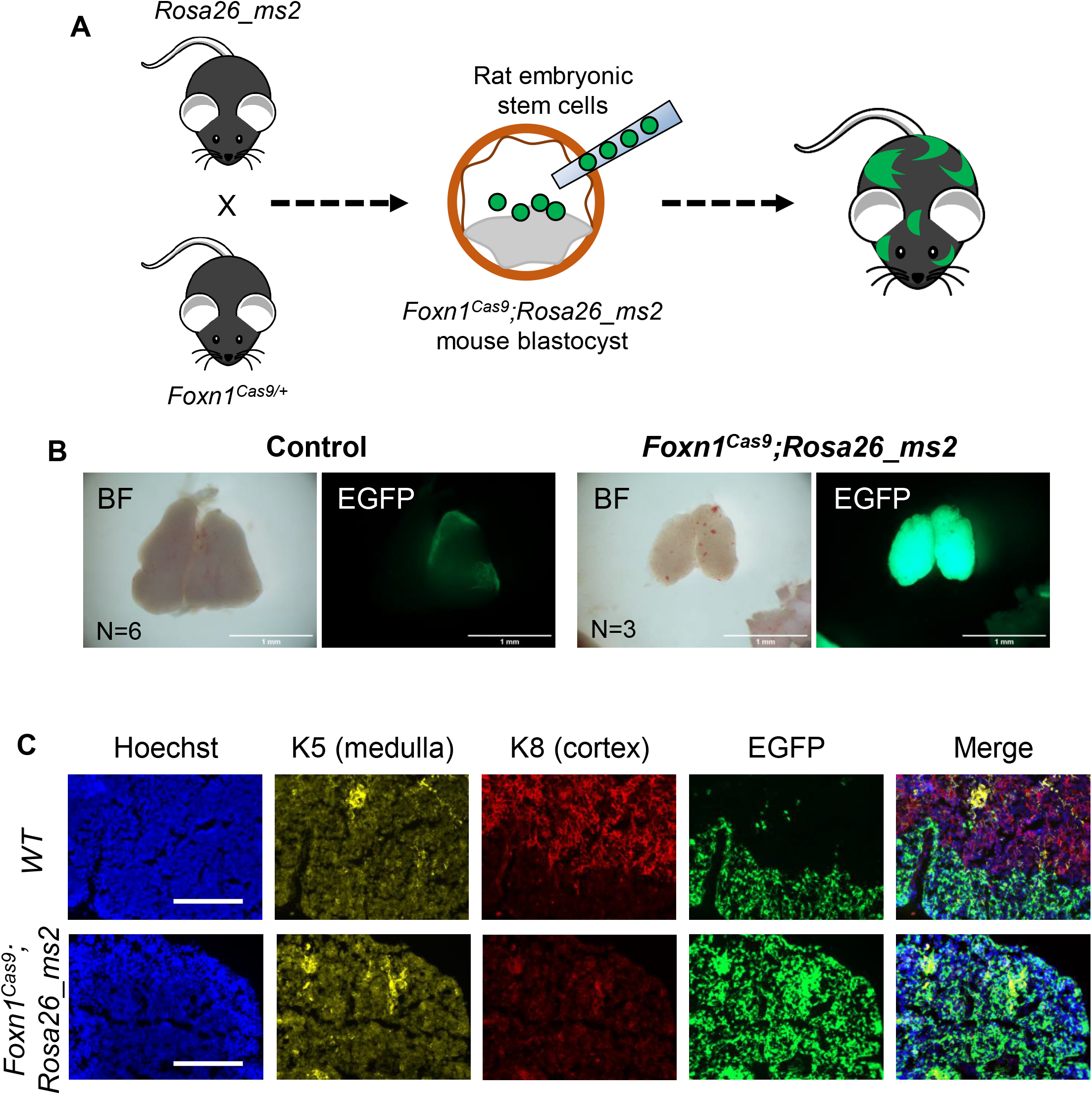
Rat thymus formation using rat ESCs and *Foxn1^Cas9^*; *Rosa26_ms* embryos by blastocyst complementation. (A) Scheme of generation of rat thymus using The Cas9-sgRNAms system (B) Thymus in *Foxn1^Cas9^*; *Rosa26_ms2* mouse host with rat ESC chimera at E18 (right). EGFP signals indicate the rat-derived cells. Wild-type littermates, *Foxn1^Cas9^*, or *Rosa26_ms2* mouse hosts with rat ESC chimeras were used as controls (left). BF means bright field. (C) Thymus sections in interspecies chimeras. Anti-K5 antibody was used to detected mouse and rat medullary thymic epithelial cells and anti-K8 antibody for mouse cortical thymic epithelial cells but not rat cortical thymic epithelial cells. The EGFP signal indicates rat thymic epithelial cells in the interspecies chimera. Hoechst indicates the nucleus. The bars show 1 mm in B and 200 µm in C.

## Discussion

### Cellular defects and athymic phenotype caused by the Cas9-sgRNA^ms^ system

The in vitro and in vivo data showed that the designed Cas9-sgRNA^ms^ system induced cell ablation through DNA DSB induction, cell apoptosis, and cell proliferation defects, and eventually led to complete ablation of the thymus. An inducible Cas9-EGFP HEK293T cell line experiment showed that Cas9-sgRNA^ms^ expression generated high levels of γH2AX and significantly increased the percentage of BrdU-negative cells. This indicated that the Cas9-sgRNA^ms^ system induced the accumulation of DNA DSB and prevented cell proliferation. Furthermore, almost all ms1 and ms2 cells showed changes in nuclear morphology following Cas9 expression (Figures 2A and 3A). In vitro data showed only a low percentage of apoptotic cells after Cas9 expression; therefore, it is possible that more cell death occurred at a later stage in the Cas9-sgRNA^ms^ system. Alternatively, they may induce cellular senescence because there are some reports that nuclear morphology (such as giant nuclei and multinuclei) and cell cycle arrest can be a biomarker of senescent cells^24–26^. Some reports show that senescent^27, 28^ and apoptotic cells^29, 30^ can be phagocytosed in vivo. Foxn1-expressed thymic epithelium may have been removed by phagocytosis after working with the Cas9-sgRNA^ms^ system.

### Skin phenotype in the *Foxn1^Cas9^*; *Rosa26_ms* mouse model

*Foxn1^nu/nu^* mice exhibit a hairless and athymic phenotype^19, 20^. However, only the thymus was affected in the *Foxn1^Cas^*^9^; *Rosa26_ms* mouse model, and not the hair phenotype. Foxn1 is also expressed in developing keratinocytes in the skin, and plays a role in hair follicle and epidermal development^31, 32^. Foxn1 expression stimulates keratinocyte growth in neighboring cells via paracrine mechanisms^32^. It is possible that even a small number of Foxn1-expressing cells remaining in the skin can maintain normal growth and differentiation of hair follicles and keratinocytes. Therefore, there may be incomplete or partial ablation of Foxn1-expressing cells in *Foxn1^Cas^*^9^; *Rosa26_ms* mouse skin.

### Contribution of rat ESCs to the thymus in the interspecies chimera

The thymus derived from rESC showed a high contribution (strong EGFP signal) to *Foxn1^Cas^*^9^; *Rosa26_ms* mouse hosts (Figure 5). The same observation was consistent with previous reports of the generation of a rat thymus in a nude mouse model^2^. The entire thymus section lacked mouse cTECs in the chimera at E18.5. However, the absence of mouse mTECs could not be proved since the marker of mTEC (anti-Keratin 5 antibody) recognized both mouse and rat Keratin 5 (Figure 5C). A study on embryonic thymus development tracking the expression of β5t (a proteasome subunit specifically expressed in cTEC but not in mTEC) showed that almost all Aire+ mTECs had a history of β5t expression^33^. Other studies show that mTECs differentiate from cells that previously expressed cTEC markers such as CD205^34^ and IL7^35^. These reports suggested that thymic progenitor cells first differentiate into cTECs and then into mTEC to form a mature thymus. Furthermore, it is unlikely that mouse cTECs preferentially differentiated into mTECs since mouse cTECs were extensively detected in the interspecies chimeric thymus (Figure 5C). Subsequently, mouse cTEC were not detected in the thymus produced with *Foxn1^Cas^*^9^; *Rosa26_ms* mouse and rat ESCs. Taken together, these results suggest that most of the thymic epithelium in the thymus produced from *Foxn1^Cas9^*; *Rosa26_ms* mouse and rat ESCs was derived from rESCs.

### Advantages of the Cas9-sgRNA^ms^ system

Cas9 was expressed under organ-specific conditions to obtain an organ-deficient animal model with constitutive expression of sgRNA^ms^. This could be potentially applied to generate animals with deficiencies in different organs and is hypothesized to ablate actively proliferating progenitor cells. The sequence of sgRNA^ms^ designed in this study has many targeting sites in the genome of several species; thus, it can be applied to generate organ-deficient mouse models and larger animal models (such as pigs) to procure human organs. The efficiency of the Cas9-sgRNA^ms^ system with the gene KO method to generate organ-deficient animal models increased from one-quarter to one-half when the Cas9-sgRNA^ms^ system was used. This can be achieved by crossing animals with a heterozygous Cas9 knock-in with another animal with a homozygous *Rosa26_ms*. This system remains an issue since the induction of the organ-deficient phenotype depends on the developmental mechanisms of the target organ. However, this system has a different mechanism of induction of cell ablation compared to the KO animal model.

### Limitations of the study

The Cas9-sgRNA^ms^ system induced cell ablation in vitro; however, the function of this system differed in different tissues and organs. Hence, the downstream response of the Cas9-sgRNA^ms^ system differs depending on the cell type and is limited to the generation of organ-deficient animal models.

## Author contributions

L.JJY. and A.I. designed experiments and performed most experiments, assisted by T.K., who provided HEK293T cell line and discussed this experimental plan, and S.Y., who performed to help all of this experiment. L.JJY. and A.I. wrote the manuscript and all authors discussed the results and commented on the manuscript.

## Acknowledgments

This work was supported by JSPS KAKENHI Grant Number 21K19267 to A.I, KAC 40th Anniversary Research Grant to A.I, and The NOVARTIS Foundation (Japan) for the Promotion of Science Grant to A.I. We appreciate to the animal experimental facility stuff for supporting to perform this study. We would like to thank to Dr. Ikawa to provide pGAG-EGxxFP plasmid and rG104 ESC line, and Dr. Yusa to provide pPB-CAG.OSKML-puDtk and pCMV-hyPBase plasmid. We would like to thank Editage (www.editage.com) for English language editing.

## Method

### Cell lines

HEK293T was cultivated in Feeder Medium, which contained 10% FBS (biosera, S00HA 10004), DMEM (Nacalai Tesque, 08459-64), L-glutamine (Thermo Fihser, 25030081), sodium private (Thermo Fihser, 11360070), Non-Essential Amino Acids (Thermo Fihser, 11140050), and penicillin-streptomycin (Thermo Fihser, 15140122). Mouse ESC line was cultivated in N2B27-a2i medium^36, 37^. Rat ESC line was cultivated in N2B27-2i medium^38, 39^.

#### Animals

Animal handling, breeding, and all experimental procedures were conducted under specific-pathogen-free (SPF) conditions at the Nara Institute of Science and Technology (NAIST), according to the guidelines of “Regulations and By-Laws of Animal Experimentation at the Nara Institute for Science and Technology”, and were approved by the Animal experimental Committee at the Nara Institute of Science and Technology (the approval no. 1639 and no. 2103). Study of the animal experiments were carried out in compliance with the ARRIVE guidelines^40^. ICR, C57BL6/J, and B6D2F1 mice used in this study were purchased from SLC, Japan, and maintained in the NAIST Animal Facility. The mouse is maintained under 12 hours light/ 12 hours dark cycle (light on 08:00 AM and light off 08:00 PM). Female ages are 8–12-week-old for generating chimeras and over the 8-week-old for breeding. Male vasectomized mice are prepared by cutting their vas deferens. Food and water are available ad libitum.

#### Plasmid constructions

pSpCas9(BB)-2A-Puro (PX459) V2.0 was purchased from Addgene (plasmid #62988)^41^. sgRNAs^ms^ and *p53* targeting sequences were ligated at the BbsI site of PX459 to produce PX459-sgRNA^ms^^1^ and PX459-sgRNA^ms^^2^. The EGFP cassette was replaced the puromycin resistance gene (Puro^r^) on the C-terminus of Cas9 in PX459 to produce pCas9-T2A-EGFP-sgRNA^ms1^ and pCas9-T2A-EGFP-sgRNA^ms2^.

sgRNAs^ms^ targeting sequences with PAM were inserted in multi-cloning-site of pCAG-EGxxFP (kindly provided by Dr. M Ikawa at Osaka University (Addgene plasmid #50716)).

pPB-TetOPuro-Cas9-EGFP plasmid for in vitro assay was produced from a Tet-One^TM^ doxycycline (Dox)-inducible system (Cat no. 634303, Takara Bio Japan) with some modifications. hPGK promoter was replaced to CAG promoter, *T2A-Puro^r^* cassette was inserted into the C-terminus of Tet-On 3G (*TetOn3G-T2A-Puro^r^*), and *Cas9-T2A-EGFP* cassette was inserted under TRE3Gs promoter. This construction was inserted between two inverted terminal repeats (ITRs) of PB5 and PB3 by replacing the CAG.OKSML-puDtk of pPB-CAG.OKSML-puDtk (kindly provided by Dr. K Yusa at Sanger Institute)^42^ to produce pPB-TetOPuro-Cas9-EGFP (Figure S2A).

pHyg-sgRNA^ms^ plasmids were produced using PX459 and pcDNA4-TO-Hygromycin-mVenus (Addgene plasmid #44099). *SV40 promoter-Hyg^r^-SV40 poly (A) signal* cassette in pcDNA4-TO-Hygromycin-mVenus was inserted and replaced to *Cbh promoter-Cas9-T2A-Puro^r^-bGH poly (A) signal* in PX459.

Targeting vectors for knock-in at *Foxn1* and *Rosa26* were constructed on the pLSODN-4D (Biodynamics Laboratory Inc.) backbone, consisting of a gene of interest flanked by left and right homology arms. The primers used for homologous arm construction are listed in Supplementary Table S1. Knock-in strategies were shown in Figure S7.

CRISPR/Cpf1 system was used for knock-in of hU6-sgRNA^ms^ cassette and Cas9 cassette at *Rosa26* and *Foxn1*. The crRNA used for Cpf1 recognition sites were ligated to BsmBI site of Cpf1-expressing vector, pTE4398 (Addgene plasmid #74042) ^43^. All Cpf1-crRNA target sequences were listed in Supplementary Table S2.

The number of sgRNAs^ms^ targeting sites, and targeting sites for *p53*, *Foxn1* and *Rosa26* were defined using CRISPRdirect software^44^

#### Generation of the Dox-inducible *Cas9-sgRNA^ms^* HEK293T cell lines

To generate the Dox-inducible *Cas9-sgRNA^ms^* HEK293T cell lines, pPB-TetOPuro-Cas9-EGFP, and pCMV-hyPBase plasmid (kindly provided by Dr. K Yusa at Sanger Institute) were co-transfected using PEI-MAX (Polysciences) into HEK293T cells and selected with puromycin (0.5µg/mL) for 7 days to establish cell lines (Figures S2B, C). Subsequently, pHyg-sgRNA^ms^ were transfected into the parental Dox-inducible Cas9-EGFP HEK293T cell line, then the Dox-inducible *Cas9-sgRNA^ms^*HEK293T cell lines were established through hygromycin B (300µg/mL) selection for 7 days.

#### Generation of *p53-KO* mouse ESC lines

Mouse ESCs, mF1-05 from 129X1/B6J F1^37^, were transfected with the sgRNA set for mouse *p53* knockout and selected using 1µg/mL puromycin for 3 days. PCR-based genotyping and karyotyping were conducted to establish *p53 ^−/−^* cell lines. sgRNAs target sites and PCR primers were listed in Supplementary Tables S2 and S3, respectively.

#### Establishment of *Foxn1^Cas9^* and *Rosa26_ms* mouse lines

Both the Cpf1-expressing plasmid and targeting vector for knock-in were transfected into mouse ESCs (mF1-05) using Lipofectamine 3000 (Thermo Fisher). G418 (150µg/mL) selection was conducted for 3 days in a2i/L media^36^, colonies were picked at ten days after transfection, and screened for knock-in through genotyping and sequencing. All primers used for genotyping are listed in Supplementary Table S3.

Established mESC lines were used to generate chimeric animals. Female ICR mice aged 8– 10 weeks were administered with 0.1 mL of CARD HyperOva® (KYUDO COMPANY) through intraperitoneal injection. After 48 h, hCG (7.5 IU) was administered through intraperitoneal injection. The superovulated female ICR mice were mated with male ICR mice. The presence of a vaginal plug the following morning indicated the occurrence of coitus and was defined as E0.5. E1.5 embryos were collected by flushing M2 media (Sigma) through the oviducts. Embryos were cultured at 37°C, 5% CO_2_ condition in KSOM media^45^ until use.

Cultured embryos at the 8-cell stage were injected with mESCs using a piezo-micromanipulator (PMAS-CT150, Prime Tech LTD) under a light microscope (DM-IRC-Leica). Each embryo was injected with six to eight mESCs in M2 medium. The injected embryos were cultured until the blastocyst stage (E3.5) in KSOM medium and transferred to the uterus of E2.5 pseudopregnant mice under anesthesia. Chimeras were recovered by natural delivery or Caesarean section on E19.5. To generate F1 mouse lines, male chimeras were mated with C57BL6/J females or B6D2F1 females. The genotype of F1 mice was identified by PCR and sequencing.

#### Generation of the interspecies chimera for blastocyst complementation

Rat ESCs tagged with EGFP, rG104 from Wistar/F344 F1 (kindly provided by Dr. Ikawa at Osaka University)^39^, were injected into mouse blastocysts that were obtained from super ovulated *Rosa26_ms* females crossed with *Foxn1^Cas9^*; *Rosa26_ms* males. Embryo collection and transferring to pseudo pregnant female mice were performed the same method as generating chimera. Generated interspecies chimeras were identified by EGFP signal derived from rat cells under a fluorescent microscope (MZFL III, Leica). The mouse host genotype of chimera was identified by PCR.

#### Flow cytometry analysis

HEK293T cells and mESCs were stained Alexa Fluor 647-conjugated annexin V, 1:50 (BioLegend) and propidium iodide (1µg/mL) in 1x annexin V binding buffer (10 mM HEPES, 140 mM NaCl, 2.5 mM CaCl_2_) on ice for 30 min. Samples were resuspended in 500µL of 1x annexin V binding buffer and filtered through a 37 µm mesh prior to cytometry analysis.

Splenocytes were collected from P9 mice by mashing the spleen between glass slides and chilling it in PBS(-). Splenocytes were stained with APC anti-mouse CD45 (1:50, Biolegend 100516) and PE anti-mouse CD3ε (1:50, Biolegend 100308) in 50 µL of 0.1% BSA/PBS at 4°C for 30 min. The samples were resuspended in 500µL of 0.1% BSA/PBS and filtered through a 37 µm mesh prior to flow cytometry analysis.

Flow cytometry was conducted using a BD Accuri C6 flow cytometer (BD Biosciences), and data were analyzed using FlowJo software (Becton Dickinson).

#### MTT assay

Inducible Cas9-sgRNA^ms^ HEK293T cells were seeded at a density of 5 × 10^3^ cells/well in 96 well plates coated with 0.2% Matrigel in the basal media. The cells were induced with doxycycline (1.5 µg/mL) for 72 h, and an MTT cell count kit (Nacalai Tesque, 23506-80) was used to perform the MTT assay. Cells were treated with 10 µL of MTT (3-(4,5-dimethylthiazol-2-yl)-2,5-diphenyltetrazolium bromide) tetrazolium solution and incubated for 3 h at 37°C, followed by adding 100 µL of solubilization solution and overnight incubation at 37°C. The absorbance at 595 nm was measured with a reference reading at 620 nm using a multi-mode plate reader (Berthold, TriStar LB942). The cell proliferation rate was calculated using the following formula:

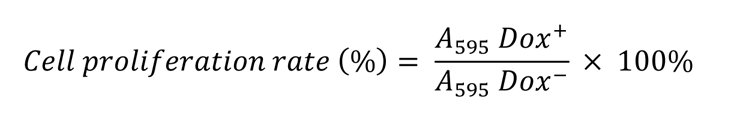

#### DNA synthesis assay

Doxycycline-inducible Cas9-sgRNA^ms^ HEK293T cells were cultured in 6-well plates and attached to glass coverslips. After 48 h of doxycycline induction, BrdU was added to the culture at a final concentration of 10 µM and further incubated for another 24 h. The cells were briefly washed in cold phosphate-buffered saline (PBS (-)) (Nacalai). 1mL 1x cytofix/cytoperm buffer (3.6% PFA and 0.2% saponin) in PBS was added to the cells and incubated at room temperature for 30 min. Buffer was removed and cells were washed once with 1 mL of wash buffer/PBS (5% FBS and 0.2% saponin). 1 mL of cytoperm plus/PBS buffer (5% BlockAce (KAC Co., Ltd.) solution, and 0.5% Triton-X 100) was then added and incubated on ice for 10 min. The buffer was then removed, and added with 1 mL of 1x cytoperm/cytofix buffer was added and incubated at room temperature for 5 min. The buffer was removed and washed with 1mL of wash buffer. Coverslips were gently removed from the 6-well plates with fine-tipped forceps and cells were treated with 50µL of 0.3 mg/mL DNaseI in PBS at 37 °C for 1 h. The cells were washed twice with the wash buffer at room temperature. Anti-BrdU antibody (1:200, Oxford Biotechnology Ltd., OB0030) was reacted on ice for 30 min. The cells were washed in wash buffer three times, Alexa Fluor 555 conjugated anti-rat IgG(H+L) antibody (Thermo Fisher, A21434) was reacted on ice for 20 min. After washing in wash buffer three times, the nuclei were stained with Hoechst33342 (1:1000; KV072, Wako) for 3 min before imaging.

#### Alkaline phosphatase (ALP) staining

ALP staining was performed using a Histofine alkaline phosphatase staining kit (Nichirei Bioscience, Japan) according to the manufacturer’s protocol. Cultured cells were washed twice in PBS and fixed by 4% paraformaldehyde (PFA) for 5 min. Freshly prepared ALP staining reagent was added and reacted at room temperature for 30 min. The cells were washed with double-distilled water prior to imaging with a digital camera (COOLPIX P7100, Nikon) or observation under a stereomicroscope.

#### Immunostaining

Cells attached to a glass coverslip or tissue section in O.C.T. compound (Sakura Finetek) were used for immunostaining. Tissue sections were prepared at 10 µm thickness using a cryostat (NX70, Leica) and dried at 37 °C for 10 min. The samples were washed twice in 2mL PBS (-) and fixed in 2% paraformaldehyde (Nacalai Tesque) for 5 min. The samples were then dehydrated with acetone (Nacalai Tesque) for 5 min. Permeabilization was performed in 50 µL 0.5% Triton X-100 (Nacalai Tesque) for 5 min. Samples were then blocked with 10% goat serum (143-06561, Wako) in BlockAce solution (KAC Co., Ltd.) at room temperature for 1 h, followed by washing in 0.1% BSA/PBS (Sigma) twice for 5 min each. Primary antibody (for DNA DSBs analysis, anti-γH2AX, 1:50, Biolegend, 613402; for immunohistochemistry of thymus, anti-K8, 1:50, Biolegend, 904804, and anti-K5, 1:50, Biolegend, 905504) was incubated at 4 °C overnight under humid conditions. The secondary antibody (Alexa Fluor 555 conjugated anti-mouse IgG, Thermo Fisher, A21425; Alexa Fluor 647 conjugated anti-rabbit IgG, Thermo Fisher, A21246) was incubated at room temperature for 1 h. Nuclei were stained with Hoechst33342 (1:1000; KV072, Wako) for 3 minutes before imaging with a fluorescent microscope (EVOS® FL, Invitrogen). Intensity of γH2AX signals were accessed using ImageJ software^46^.

#### Statistical analysis

Statistical tests were performed using Prism 9 (GraphPad) or EZR software^47^. The confidence interval was set at 95%, and a *p-*value less than 0.05 indicates a significant difference between the compared datasets.

---

**Figure S1.**
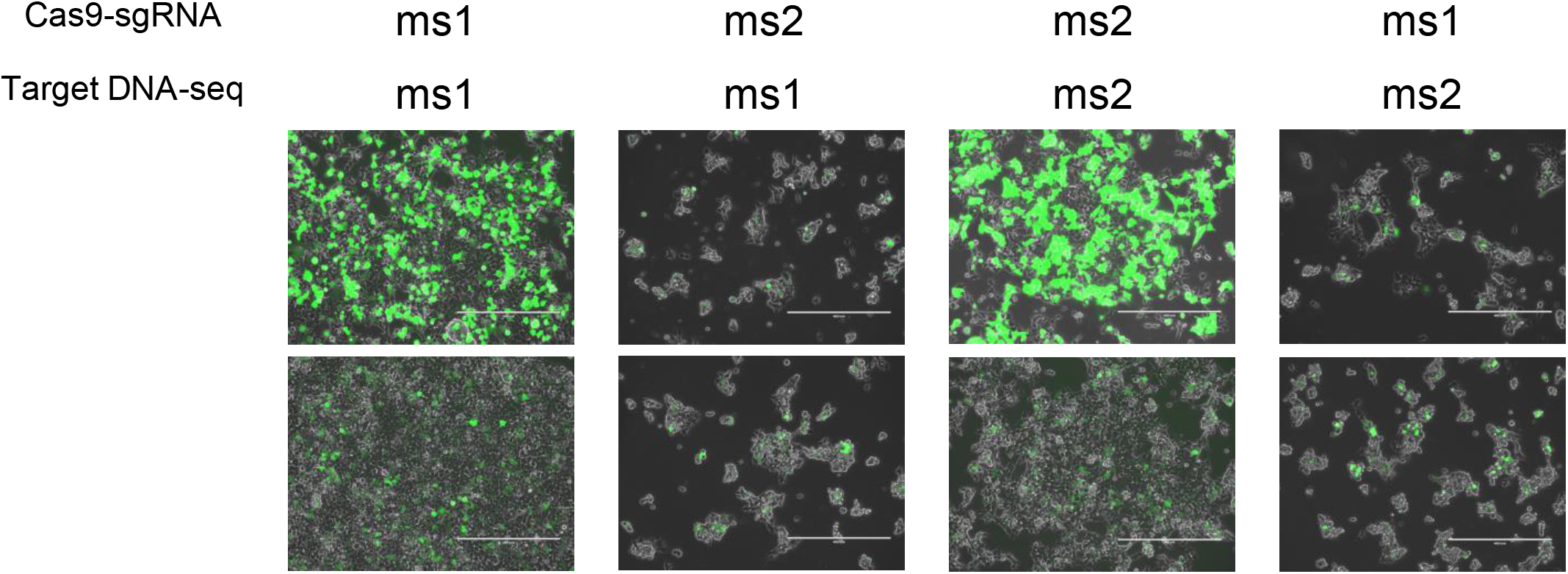
Validation of sgRNAms sequence specificity in HEK293T. HEK293T cells were transfected with Cas9-sgRNA^ms^ and matching sgRNA_ms-target or non-matching sgRNA-target vectors. Green signals indicate that the sgRNA target in pCAG-EGxxFP was cleaved by the Cas9-sgRNA^ms^system; subsequently, it occurred as a homology-dependent repair and then reconstituted the EGFP expression cassette. Scale bars, 400µm.

**Figure S2.**
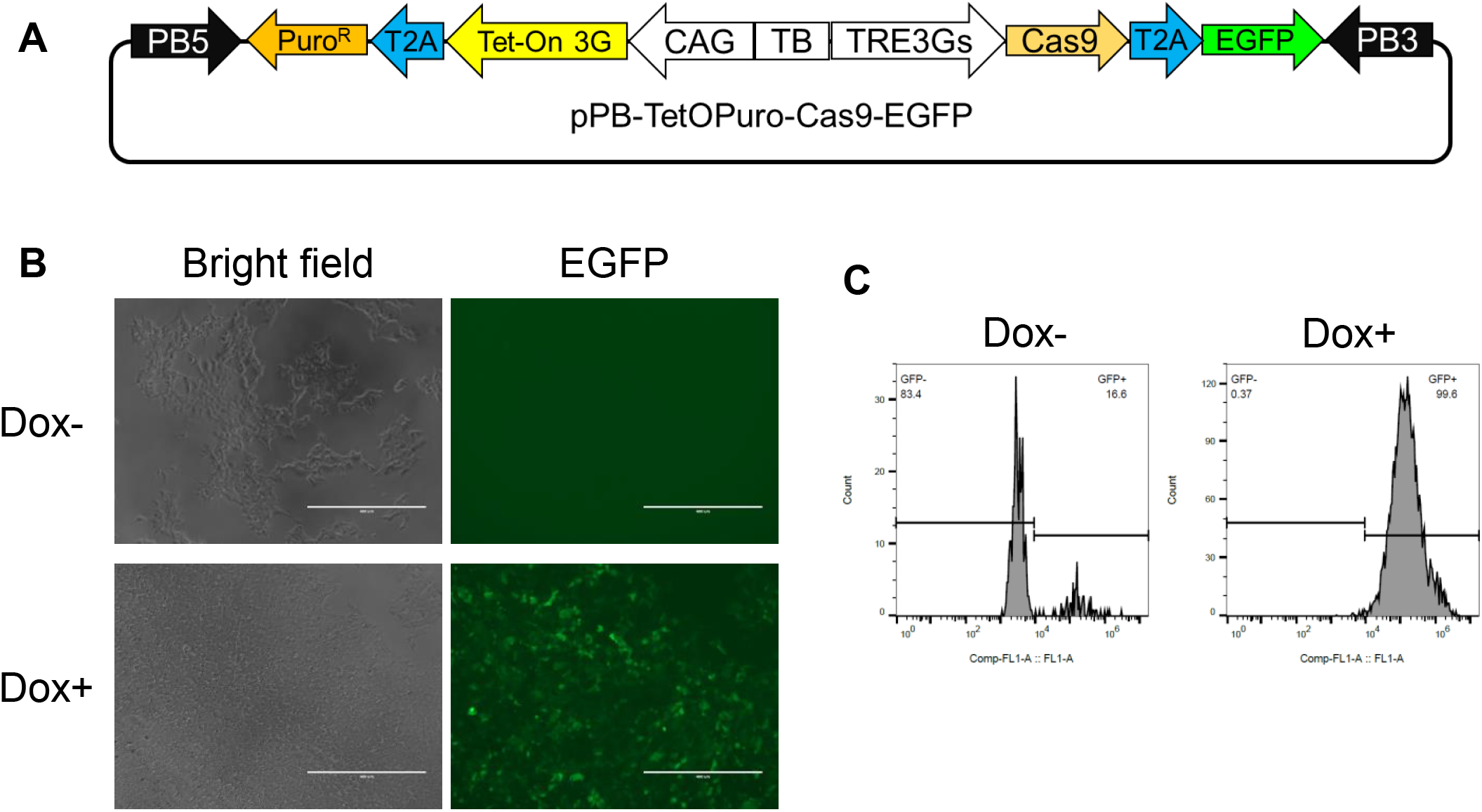
Doxycycline inducible Cas9 in HEK293T. (A) Construction of a doxycycline (Dox)-inducible Cas9-T2A-EGFP vector. (B) and (C) monitoring Cas9 expression by EGFP signaling after Dox induction. Dox-indicates no Dox in the culture medium, whereas Dox+ indicates Dox in the culture medium. Bars show 400 µm in B.

**Figure S3.**
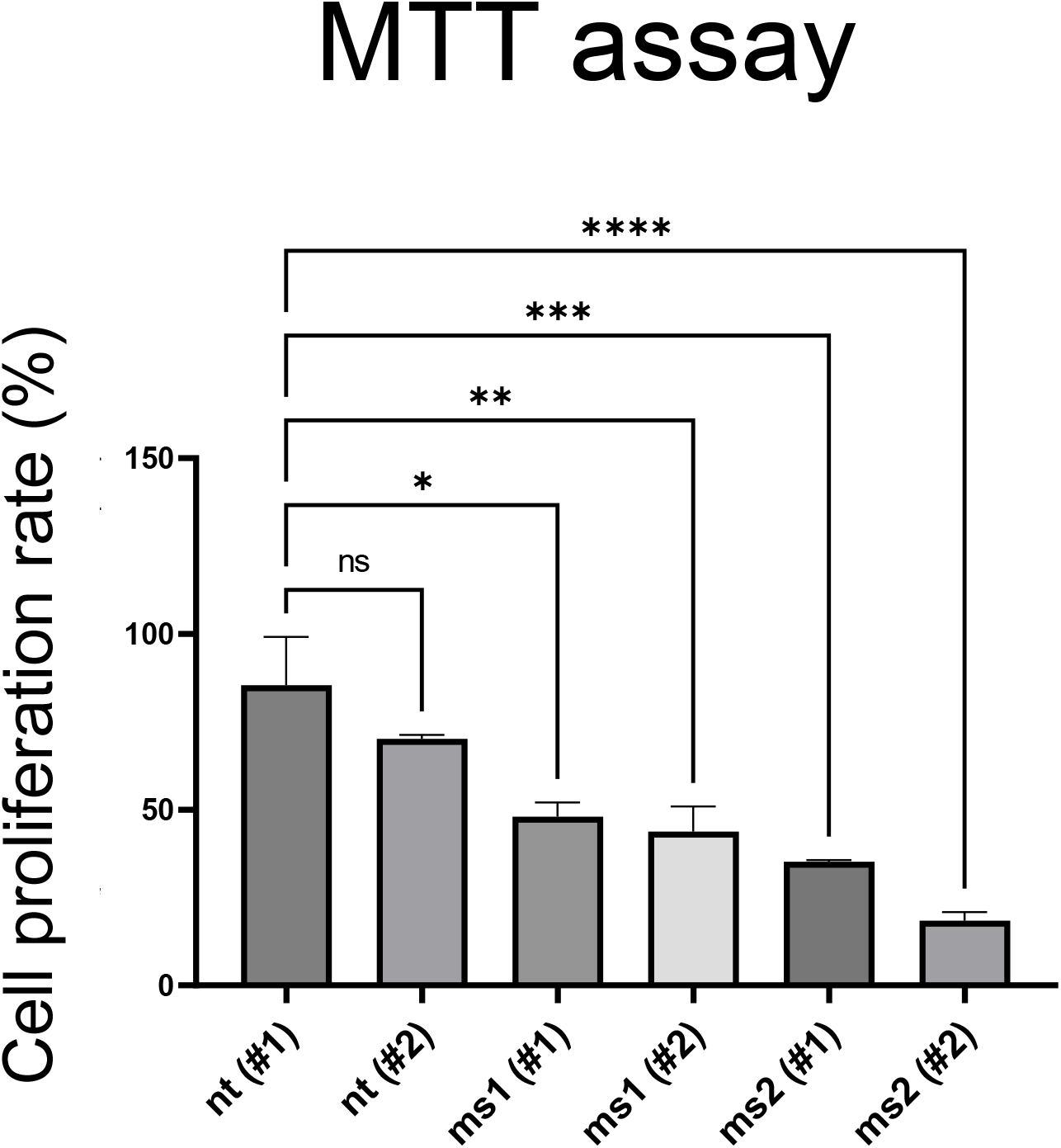
Cell proliferation assay in HEK293T using the MTT assay. Proliferation rate of HEK293T cells expressing sgRNA^ms1^ (ms1), sgRNA^ms2^ (ms2), and non-targeted sgRNA (nt) after 72 h of DOX-induced Cas9-EGFP expression. One-way ANOVA, Tukey honestly significant difference test was used for statistical analysis. *p<0.05; **p<0.01; ***p<0.001; ****p<0.0001.

**Figure S4.**
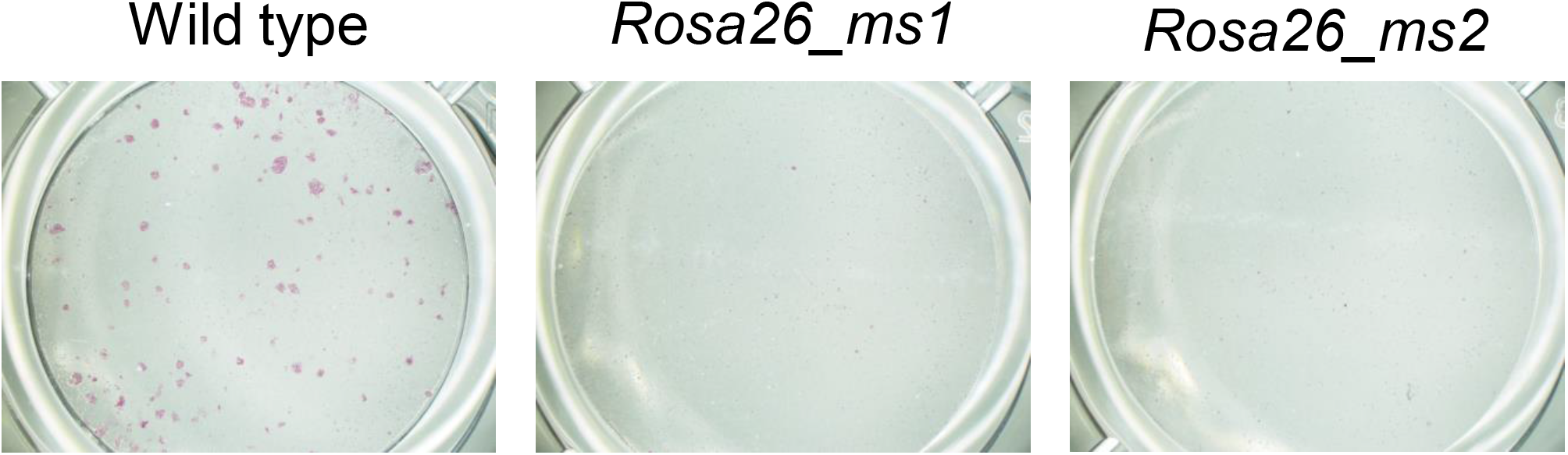
Effects of Cas9-sgRNA^ms^ system in mouse ESCs. Colony formation after transient Cas9 expression in the mouse ESC line, which has a ubiquitously expressed sgRNA^ms^ cassette knocked-in the *ROSA26* locus. Alkaline phosphatase-positive colonies are shown in light violet and represent surviving mESCs.

**Figure S5.**
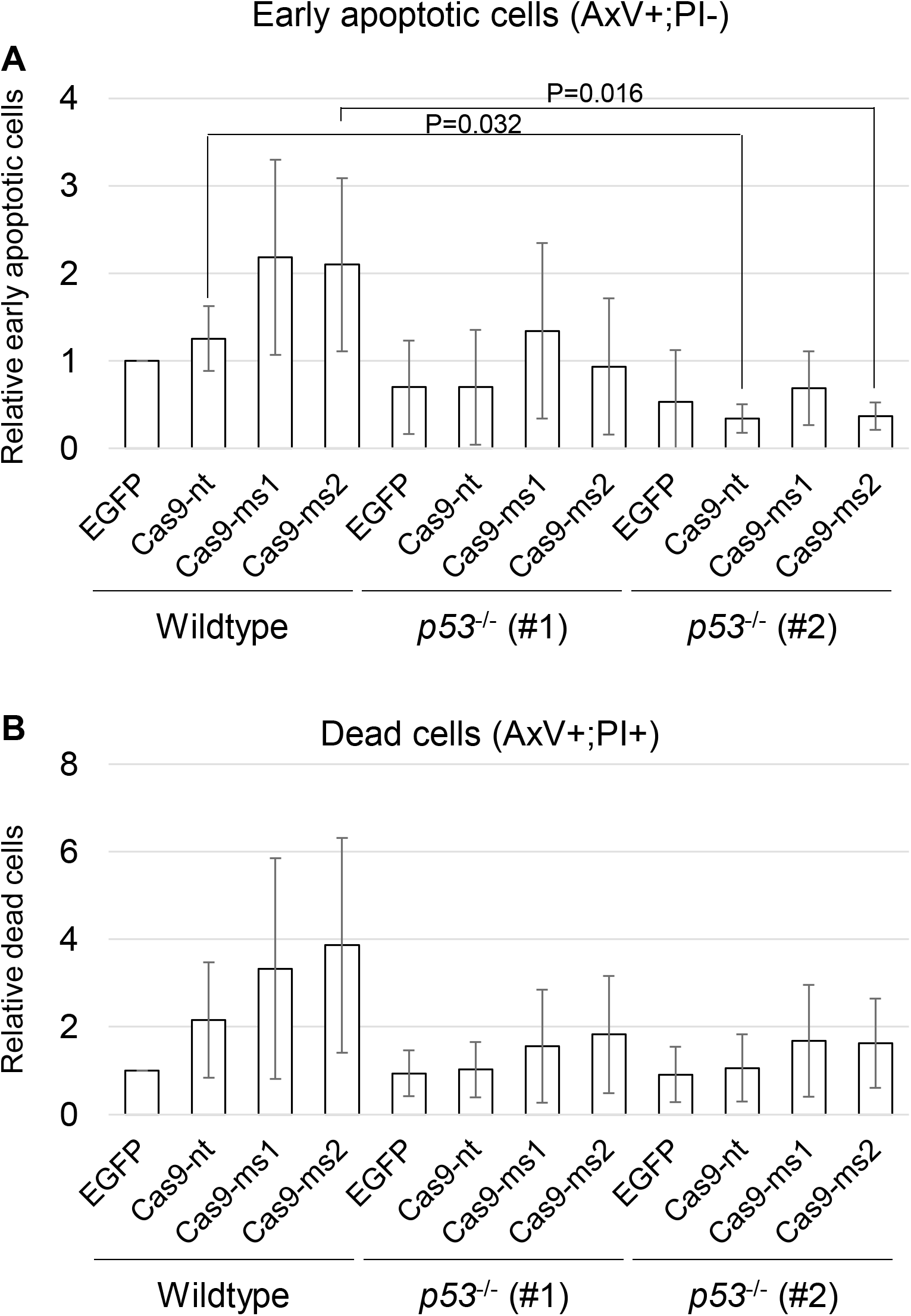
Involvement of p53 in mESCs functioning with the Cas9-sgRNA^ms^ system. Relative populations of early apoptotic cells (A) and dead cells (B) 72 h after transfection with Cas9-sgRNA^ms^ expression vectors in wildtype ESCs as control and two lines of *p53* knock-out mouse ESCs. Mean ± SEM. Two-way ANOVA was performed for the analysis of statistical differences. Significant differences were observed between the factor of ESC lines (P=8.4×10-5) in A, and both factors of vectors (P=0.016) and ESC lines (P=0.037) in B. Dunnett’s test were performed after One-way ANOVA analysis for between-group comparisons. No significant differences were found in B.

**Figure S6.**
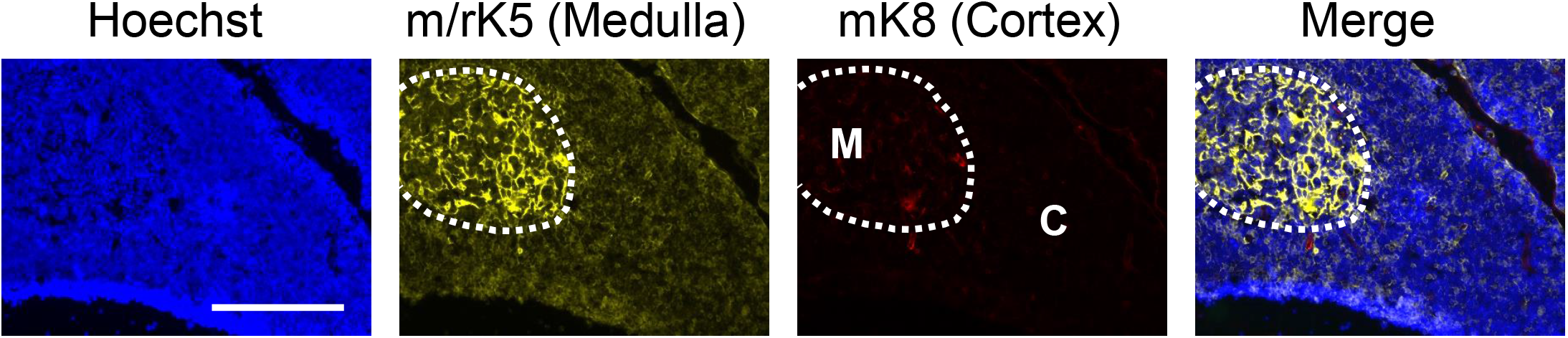
Immunohistochemistry of the 7-day-old rat thymus. The anti-K5 antibody was detected in rat medullary thymic epithelial cells, whereas the anti-K8 antibody did not recognize rat cortical thymic epithelial cells, even in 7-day-old rats. Bar shows 200 µm.

**Figure S7.**
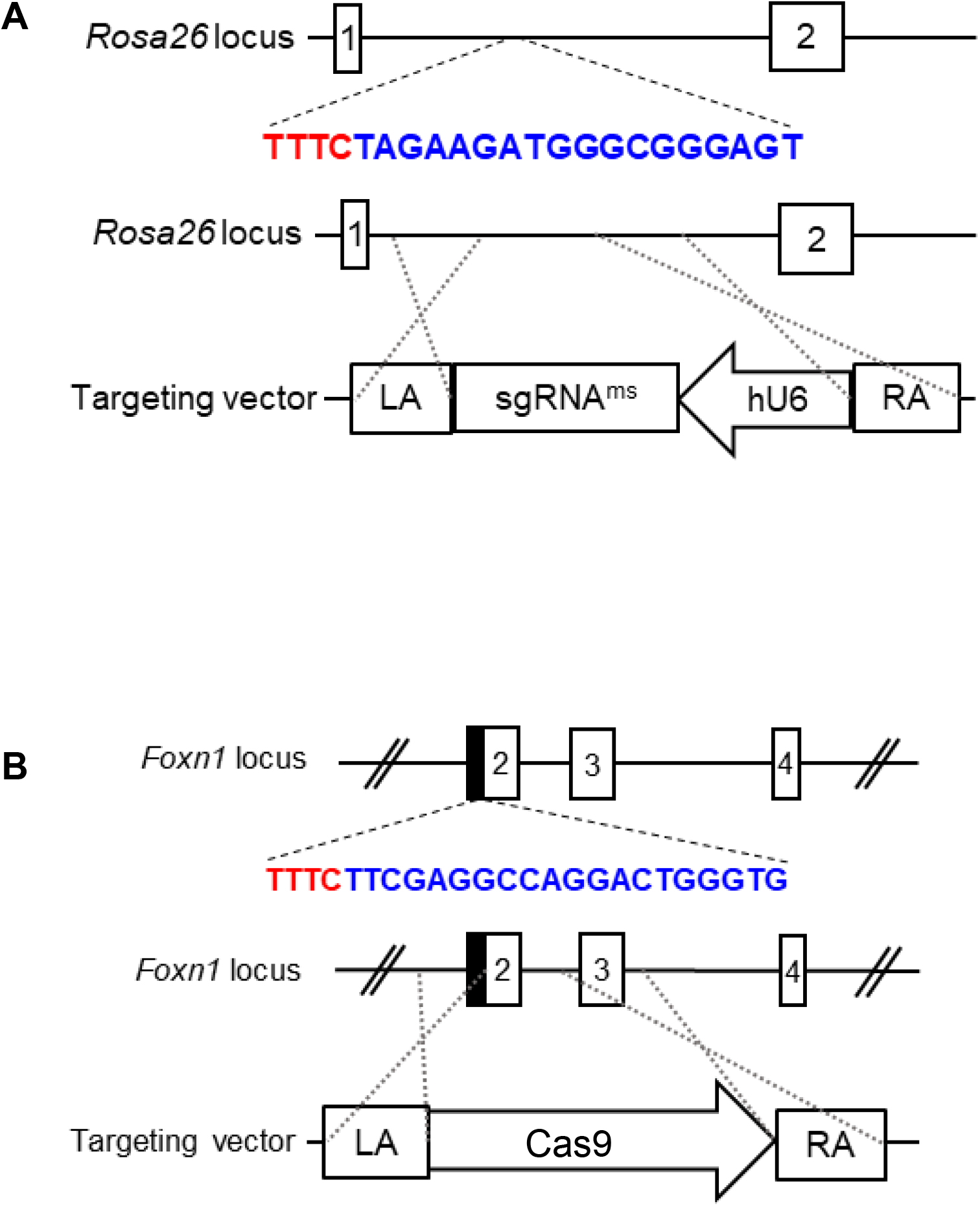
Schematic diagram of knock-in strategy at different loci. (A) *Rosa26_ms*; (B) *Foxn1^Cas9^* Red: PAM sequence of *CpfI*; Blue: crRNA sequence; LA: Left homologous arm; RA: Right homologous arm.

**Supplementary table S1.**
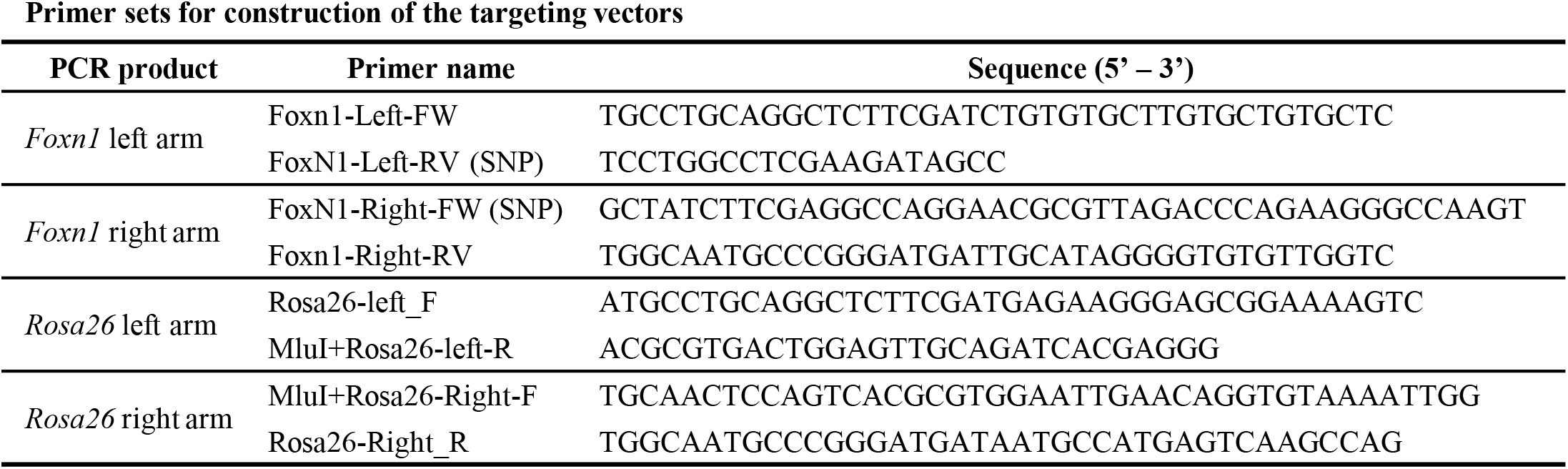
Primer sets for construction of the targeting vectors.

**Supplementary table S2.**
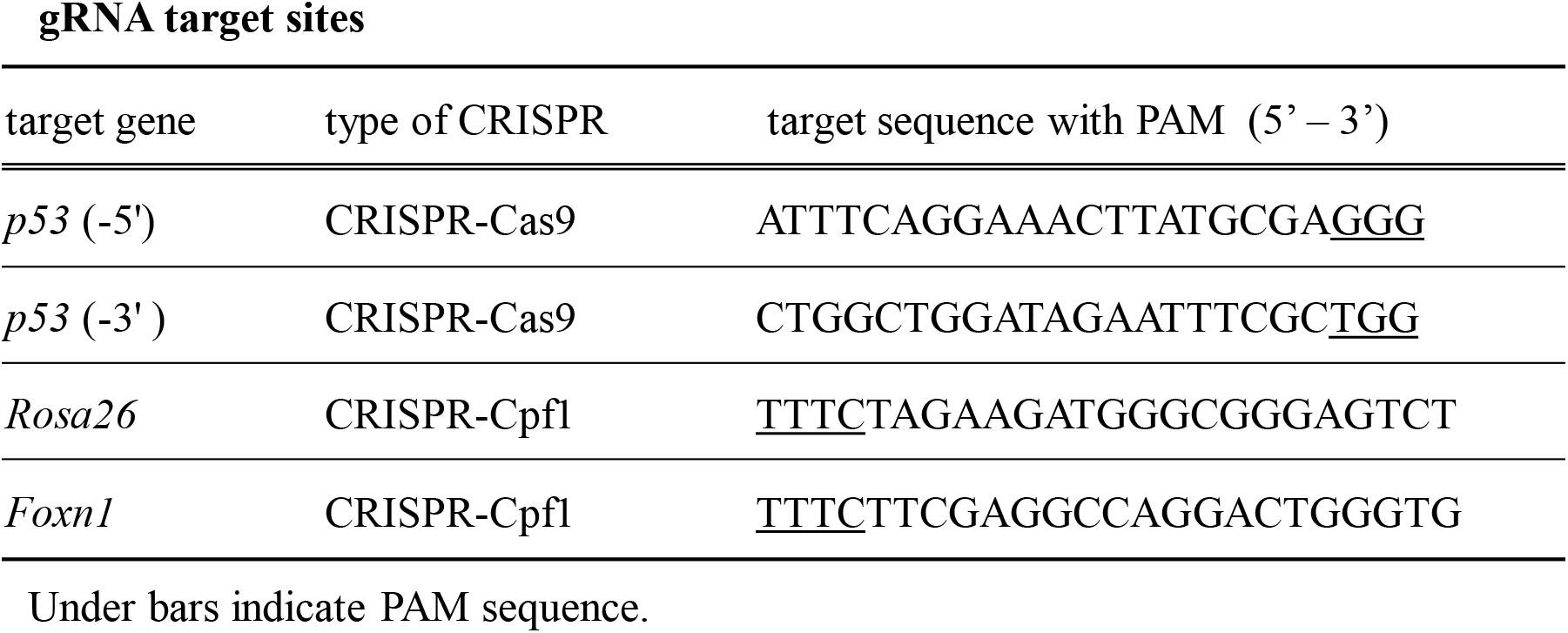
gRNA target sites.

**Supplementary table S3.**
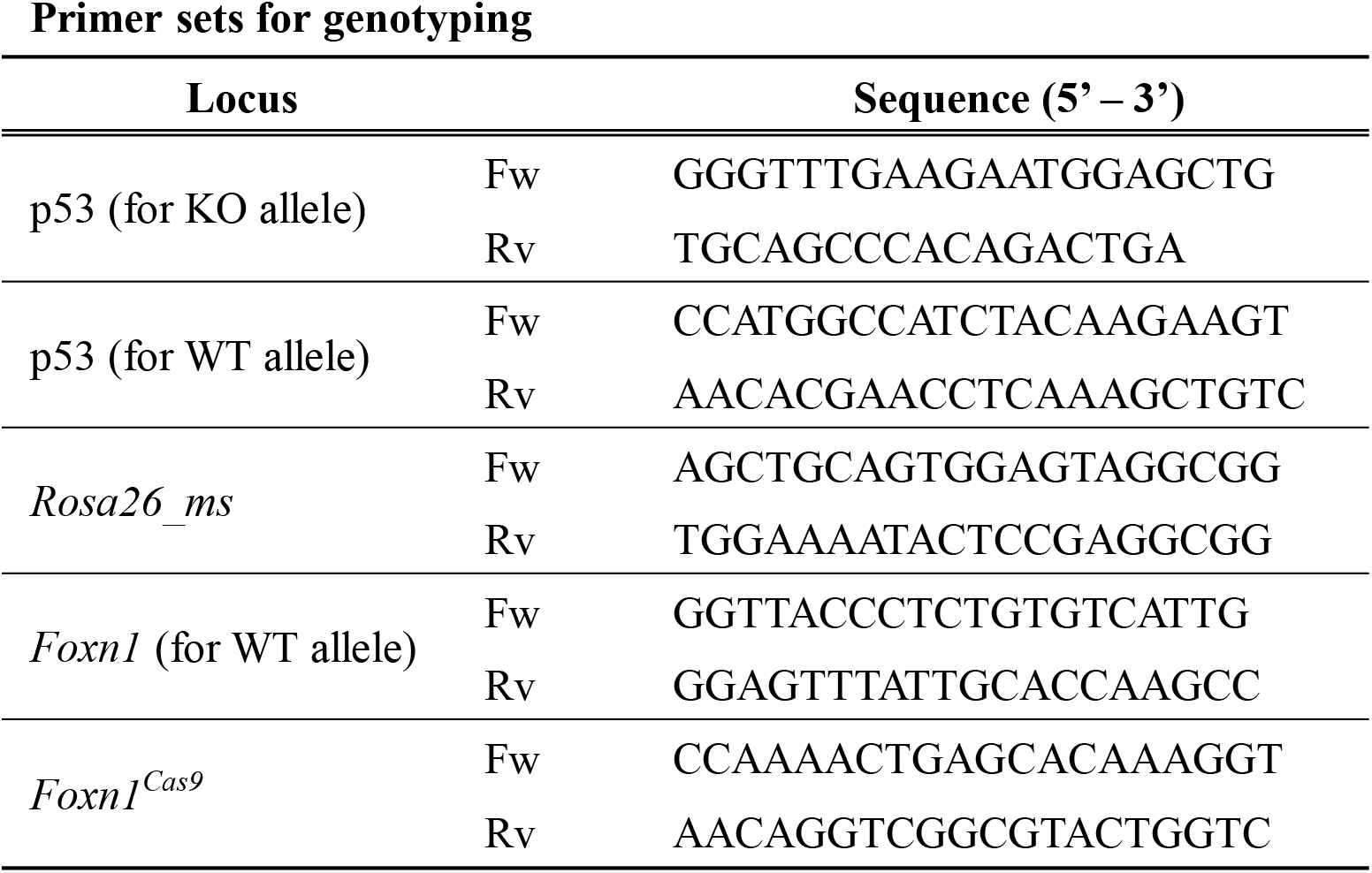
Primer sets for genotyping.

## References

1. Kobayashi T, Yamaguchi T, Hamanaka S, et al. Generation of rat pancreas in mouse by interspecific blastocyst injection of pluripotent stem cells. Cell. Sep 03 2010;142(5):787–99. doi:10.1016/j.cell.2010.07.039

2. Isotani A, Hatayama H, Kaseda K, Ikawa M, Okabe M. Formation of a thymus from rat ES cells in xenogeneic nude mouse↔rat ES chimeras. Genes Cells. Apr 2011;16(4):397–405. doi:10.1111/j.1365-2443.2011.01495.x

3. Yamaguchi T, Sato H, Kato-Itoh M, et al. Interspecies organogenesis generates autologous functional islets. Nature. Feb 09 2017;542(7640):191-196. doi:10.1038/nature21070

4. Goto T, Hara H, Sanbo M, et al. Generation of pluripotent stem cell-derived mouse kidneys in Sall1-targeted anephric rats. Nat Commun. Feb 05 2019;10(1):451. doi:10.1038/s41467-019-08394-9

5. Kobayashi T, Goto T, Oikawa M, et al. Blastocyst complementation using Prdm14-deficient rats enables efficient germline transmission and generation of functional mouse spermatids in rats. Nat Commun. Feb 26 2021;12(1):1328. doi:10.1038/s41467-021-21557-x

6. Mori M, Furuhashi K, Danielsson JA, et al. Generation of functional lungs via conditional blastocyst complementation using pluripotent stem cells. Nat Med. Nov 2019;25(11):1691–1698. doi:10.1038/s41591-019-0635-8

7. Watanabe M, Nakano K, Uchikura A, et al. Anephrogenic phenotype induced by SALL1 gene knockout in pigs. Sci Rep. May 29 2019;9(1):8016. doi:10.1038/s41598-019-44387-w

8. Kitahara A, Ran Q, Oda K, et al. Generation of Lungs by Blastocyst Complementation in Apneumic Fgf10-Deficient Mice. Cell Rep. May 12 2020;31(6):107626. doi:10.1016/j.celrep.2020.107626

9. Matsumura H, Hasuwa H, Inoue N, Ikawa M, Okabe M. Lineage-specific cell disruption in living mice by Cre-mediated expression of diphtheria toxin A chain. Biochem Biophys Res Commun. Aug 20 2004;321(2):275–9. doi:10.1016/j.bbrc.2004.06.139

10. Ivanova A, Signore M, Caro N, Greene ND, Copp AJ, Martinez-Barbera JP. In vivo genetic ablation by Cre-mediated expression of diphtheria toxin fragment A. Genesis. Nov 2005;43(3):129–35. doi:10.1002/gene.20162

11. Stanger BZ, Tanaka AJ, Melton DA. Organ size is limited by the number of embryonic progenitor cells in the pancreas but not the liver. Nature. Feb 22 2007;445(7130):886-91. doi:10.1038/nature05537

12. Saito M, Iwawaki T, Taya C, et al. Diphtheria toxin receptor-mediated conditional and targeted cell ablation in transgenic mice. Nat Biotechnol. Aug 2001;19(8):746–50. doi:10.1038/90795

13. Jinek M, Chylinski K, Fonfara I, Hauer M, Doudna JA, Charpentier E. A programmable dual-RNA-guided DNA endonuclease in adaptive bacterial immunity. Science. Aug 17 2012;337(6096):816-21. doi:10.1126/science.1225829

14. Kwon T, Ra JS, Lee S, et al. Precision targeting tumor cells using cancer-specific InDel mutations with CRISPR-Cas9. Proc Natl Acad Sci U S A. Mar 01 2022;119(9)doi:10.1073/pnas.2103532119

15. Głów D, Maire CL, Schwarze LI, Lamszus K, Fehse B. CRISPR-to-Kill (C2K)-Employing the Bacterial Immune System to Kill Cancer Cells. Cancers (Basel). Dec 15 2021;13(24)doi:10.3390/cancers13246306

16. Martinez-Lage M, Torres-Ruiz R, Puig-Serra P, et al. In vivo CRISPR/Cas9 targeting of fusion oncogenes for selective elimination of cancer cells. Nat Commun. Oct 08 2020;11(1):5060. doi:10.1038/s41467-020-18875-x

17. Bharat A, Mohanakumar T. Immune Responses to Tissue-Restricted Nonmajor Histocompatibility Complex Antigens in Allograft Rejection. J Immunol Res. 2017;2017:6312514. doi:10.1155/2017/6312514

18. Vaidya HJ, Briones Leon A, Blackburn CC. FOXN1 in thymus organogenesis and development. Eur J Immunol. Aug 2016;46(8):1826–37. doi:10.1002/eji.201545814

19. Flanagan SP. ’Nude’, a new hairless gene with pleiotropic effects in the mouse. Genet Res. Dec 1966;8(3):295–309. doi:10.1017/s0016672300010168

20. Nehls M, Pfeifer D, Schorpp M, Hedrich H, Boehm T. New member of the winged-helix protein family disrupted in mouse and rat nude mutations. Nature. Nov 03 1994;372(6501):103-7. doi:10.1038/372103a0

21. Pantelouris EM. Absence of thymus in a mouse mutant. Nature. Jan 27 1968;217(5126):370-1. doi:10.1038/217370a0

22. Mashiko D, Fujihara Y, Satouh Y, Miyata H, Isotani A, Ikawa M. Generation of mutant mice by pronuclear injection of circular plasmid expressing Cas9 and single guided RNA. Sci Rep. Nov 27 2013;3:3355. doi:10.1038/srep03355

23. Burma S, Chen BP, Murphy M, Kurimasa A, Chen DJ. ATM phosphorylates histone H2AX in response to DNA double-strand breaks. J Biol Chem. Nov 09 2001;276(45):42462–7. doi:10.1074/jbc.C100466200

24. Kumari R, Jat P. Mechanisms of Cellular Senescence: Cell Cycle Arrest and Senescence Associated Secretory Phenotype. Front Cell Dev Biol. 2021;9:645593. doi:10.3389/fcell.2021.645593

25. Di Micco R, Krizhanovsky V, Baker D, d’Adda di Fagagna F. Cellular senescence in ageing: from mechanisms to therapeutic opportunities. Nat Rev Mol Cell Biol. Feb 2021;22(2):75–95. doi:10.1038/s41580-020-00314-w

26. González-Gualda E, Baker AG, Fruk L, Muñoz-Espín D. A guide to assessing cellular senescence in vitro and in vivo. FEBS J. Jan 2021;288(1):56–80. doi:10.1111/febs.15570

27. Kale A, Sharma A, Stolzing A, Desprez PY, Campisi J. Role of immune cells in the removal of deleterious senescent cells. Immun Ageing. 2020;17:16. doi:10.1186/s12979-020-00187-9

28. Behmoaras J, Gil J. Similarities and interplay between senescent cells and macrophages. J Cell Biol. Feb 01 2021;220(2)doi:10.1083/jcb.202010162

29. Poon IK, Lucas CD, Rossi AG, Ravichandran KS. Apoptotic cell clearance: basic biology and therapeutic potential. Nat Rev Immunol. Mar 2014;14(3):166–80. doi:10.1038/nri3607

30. Elliott MR, Ravichandran KS. The Dynamics of Apoptotic Cell Clearance. Dev Cell. Jul 25 2016;38(2):147–60. doi:10.1016/j.devcel.2016.06.029

31. Kur-Piotrowska A, Kopcewicz M, Kozak LP, Sachadyn P, Grabowska A, Gawronska-Kozak B. Neotenic phenomenon in gene expression in the skin of Foxn1-deficient (nude) mice-a projection for regenerative skin wound healing. BMC Genomics. Jan 09 2017;18(1):56. doi:10.1186/s12864-016-3401-z

32. Prowse DM, Lee D, Weiner L, et al. Ectopic expression of the nude gene induces hyperproliferation and defects in differentiation: implications for the self-renewal of cutaneous epithelia. Dev Biol. Aug 01 1999;212(1):54–67. doi:10.1006/dbio.1999.9328

33. Ohigashi I, Zuklys S, Sakata M, et al. Aire-expressing thymic medullary epithelial cells originate from β5t-expressing progenitor cells. Proc Natl Acad Sci U S A. Jun 11 2013;110(24):9885–90. doi:10.1073/pnas.1301799110

34. Baik S, Jenkinson EJ, Lane PJ, Anderson G, Jenkinson WE. Generation of both cortical and Aire(+) medullary thymic epithelial compartments from CD205(+) progenitors. Eur J Immunol. Mar 2013;43(3):589–94. doi:10.1002/eji.201243209

35. Ribeiro AR, Rodrigues PM, Meireles C, Di Santo JP, Alves NL. Thymocyte selection regulates the homeostasis of IL-7-expressing thymic cortical epithelial cells in vivo. J Immunol. Aug 01 2013;191(3):1200–9. doi:10.4049/jimmunol.1203042

36. Choi J, Huebner AJ, Clement K, et al. Prolonged Mek1/2 suppression impairs the developmental potential of embryonic stem cells. Nature. 08 2017;548(7666):219-223. doi:10.1038/nature23274

37. Kishimoto Y, Nishiura I, Hirata W, et al. A novel tissue specific alternative splicing variant mitigates phenotypes in Ets2 frame-shift mutant models. Sci Rep. Apr 2021;11(1):8297. doi:10.1038/s41598-021-87751-5

38. Li P, Tong C, Mehrian-Shai R, et al. Germline competent embryonic stem cells derived from rat blastocysts. Cell. Dec 26 2008;135(7):1299–310. doi:10.1016/j.cell.2008.12.006

39. Isotani A, Yamagata K, Okabe M, Ikawa M. Generation of Hprt-disrupted rat through mouse←rat ES chimeras. Sci Rep. Apr 2016;6:24215. doi:10.1038/srep24215

40. Kilkenny C, Browne WJ, Cuthill IC, Emerson M, Altman DG. Improving bioscience research reporting: the ARRIVE guidelines for reporting animal research. PLoS Biol. Jun 29 2010;8(6):e1000412. doi:10.1371/journal.pbio.1000412

41. Ran FA, Hsu PD, Wright J, Agarwala V, Scott DA, Zhang F. Genome engineering using the CRISPR-Cas9 system. Nat Protoc. Nov 2013;8(11):2281–2308. doi:10.1038/nprot.2013.143

42. Yusa K, Rad R, Takeda J, Bradley A. Generation of transgene-free induced pluripotent mouse stem cells by the piggyBac transposon. Nat Methods. May 2009;6(5):363–9. doi:10.1038/nmeth.1323

43. Tóth E, Weinhardt N, Bencsura P, et al. Cpf1 nucleases demonstrate robust activity to induce DNA modification by exploiting homology directed repair pathways in mammalian cells. Biol Direct. Sep 14 2016;11:46. doi:10.1186/s13062-016-0147-0

44. Naito Y, Hino K, Bono H, Ui-Tei K. CRISPRdirect: software for designing CRISPR/Cas guide RNA with reduced off-target sites. Bioinformatics. Apr 2015;31(7):1120–3. doi:10.1093/bioinformatics/btu743

45. Ho Y, Wigglesworth K, Eppig JJ, Schultz RM. Preimplantation development of mouse embryos in KSOM: augmentation by amino acids and analysis of gene expression. Mol Reprod Dev. Jun 1995;41(2):232–8. doi:10.1002/mrd.1080410214

46. Schneider CA, Rasband WS, Eliceiri KW. NIH Image to ImageJ: 25 years of image analysis. Nat Methods. Jul 2012;9(7):671–5. doi:10.1038/nmeth.2089

47. Kanda Y. Investigation of the freely available easy-to-use software ’EZR’ for medical statistics. Bone Marrow Transplant. Mar 2013;48(3):452–8. doi:10.1038/bmt.2012.244

